# Detecting expressed cancer somatic mutations from single-cell RNA sequencing data

**DOI:** 10.1101/2021.10.08.463191

**Authors:** Tianyun Zhang, Ning Shen

## Abstract

Identifying expressed somatic mutations directly from single-cell RNA sequencing (scRNA-seq) data is challenging but highly valuable. Computational methods have been attempted but no reliable methods have been reported to identify somatic mutations with high fidelity. We present RESA -- Recurrently Expressed SNV Analysis, a computational framework that identifies expressed somatic mutations from scRNA-seq data with high precision. We test RESA in multiple cancer cell line datasets, where RESA demonstrates average area under the curve (AUC) of 0.9 on independently held out test sets, and achieves average precision of 0.71 when evaluated by bulk whole exome, which is substantially higher than previous approaches. In addition, RESA detects a median of 201 mutations per cell, 50 times more than what was reported in experimental technologies with simultaneous expression and mutation profiling. Furthermore, applying RESA to scRNA-seq from a melanoma patient, we demonstrate that RESA recovers the known BRAF driver mutation of the sample and melanoma dominating mutational signatures, identifies mutation associated expression signatures, reveals nondriver perturbed and stage specific cancer hallmarks, and unveils the complex relationship between genomic and transcriptomic intratumor heterogeneity. Therefore, RESA could provide novel views in the study of intratumor heterogeneity and relate genetic alterations to transcriptional changes at single cell level.

## Introduction

Somatic mutations accumulate in cell generations and are often found to transform normal cells into cancer and promote tumor progression. Genetic heterogeneity as well as transcriptional heterogeneity in cancer are both key aspects that contribute to cancer evolution and drug resistance. However, it remains challenging to relate the transcriptional heterogeneity to genetic heterogeneity at single cell level. In this regard, identifying somatic mutations carried by the RNA at single cell level is highly valuable.

Due to the limited biological sample available in a single cell, simultaneous experimental profiling of both DNA and RNA with high coverage remains challenging. Despite the development of experimental technologies of targeted genotyping in scRNA-seq (Giustacchini *et al*., 2017; Nam *et al*., 2019; Rodriguez-Meira *et al*., 2019), the technical challenges, small number of detectable mutations and low signal-to-noise ratio have limited their application. On the other hand, the value of computational methods that identify expressed somatic mutations from scRNA-seq data is widely appreciated. However, identifying expressed somatic mutations from scRNA-seq is challenging due to both sparsity of real signal and high levels of noise and artifacts. As a result, profiling of scRNA-seq with matched bulk genotyping, e.g. whole exome sequencing (WES) or whole genome sequencing (WGS), and single cell genotyping of selected mutations, of the same sample have been applied to study intratumor heterogeneity and lineage tracing (Giustacchini *et al*., 2017; L. Wang *et al*., 2017; Campbell *et al*., 2019; Hovestadt *et al*., 2019; Petti *et al*., 2019; van Galen *et al*., 2019). Such data can be very helpful and effective in eliminating false positives, but require thoughtful design, large amounts of samples, and careful analysis, thus was not widely adopted.

Computational methods to identify expressed somatic mutations using only scRNA-seq are much more appealing in comparison, even with limited sensitivity, as such methods not only provide orthogonal insights into the intratumor heterogeneity, but can be applied for analysis in a retrospective manner as well. Mutational analysis of scRNA-seq are mostly using data generated from scRNA-seq technologies with full length library prep, e.g. SMART-seq2 (Picelli *et al*., 2013) for its higher coverage in the gene body compared to 10X genomics and Drop-seq approach, and have been applied to various biological questions including aging in human pancreas, Alzheimer’s disease, glioblastoma, lung cancer, etc (Darmanis *et al*., 2017; Enge *et al*., 2017; Liu *et al*., 2019; Mathys *et al*., 2019; Maynard *et al*., 2020). Such methods have been explored with differences in details, but are generally carried out by either combining variant callers with quality-based filtering, or stratifying from a whitelist of curated known cancer mutations. However, these methods either result in large amounts of artefact signals as reported or have not been assessed on the reliability of detection.

In this study, we explored the detectability of cancer somatic single nucleotide variants (SNVs) from scRNA-seq. We focused on scRNA-seq technologies that capture full length transcripts, e.g. SMART-seq2. We demonstrated that a number of factors like expression variability and clonal effects could impact the detectability of expressed somatic mutations. To overcome the high level of noise, we developed a computational pipeline named Recurrently Expressed SNV Analysis (RESA) that can detect expressed somatic SNVs with high precision directly from scRNA-seq data.

RESA outperforms existing methods in multiple aspects. First, the filtering and analytical processing of RESA substantially improves the precision of somatic variant detection from scRNA-seq data compared to existing methods. RESA consistently demonstrates substantially higher precision than quality-based filtering and whitelist based filtering methods across multiple independently generated datasets. Secondly, RESA expands the candidate true somatic variant set using a custom designed logistic regression classifier. This helps improve the detection sensitivity in the expected sparse scRNA-seq dataset while maintaining high precision. Most importantly, RESA allows detection of expressed mutations beyond cancer driver mutations, recapitulates mutational signatures at population level, and enables study of active biological processes carrying somatic mutations on top of expression variation in scRNA-seq.

## Results

### Sparsity and noisiness of expressed mutations in scRNA-seq data

Identifying somatic mutations directly from scRNA-seq data is challenging, due to both sparsity and noisiness. Genomic mutations that are reliably expressed and detectable through scRNA-seq data are expected to be sparse. The effect of site-specific depth variation on mutation detectability can be demonstrated by coverage comparison of multi-omic profiling. Multi-omic data for the prostate cancer cell line LNCaP was used for demonstration purpose hereafter. For example, a hotspot mutation in the TBL1XR1 gene exon in the LNCaP cell line has high sitespecific sequencing depth in both WES and bulk RNA-seq data, thus are easily detectable. However, in the scRNA-seq data of the same cell line, two of five selected single cells have few to no site-specific depth, prohibiting detection of the mutation (Fig. 1a). Indeed, of all the exonic somatic SNVs identified from WES for LNCaP, only 2%-8% of exonic SNVs per cell was detectable with sufficient site-specific depth in the corresponding scRNA-seq dataset (Fig. 1b, Methods).

**Figure 1.**
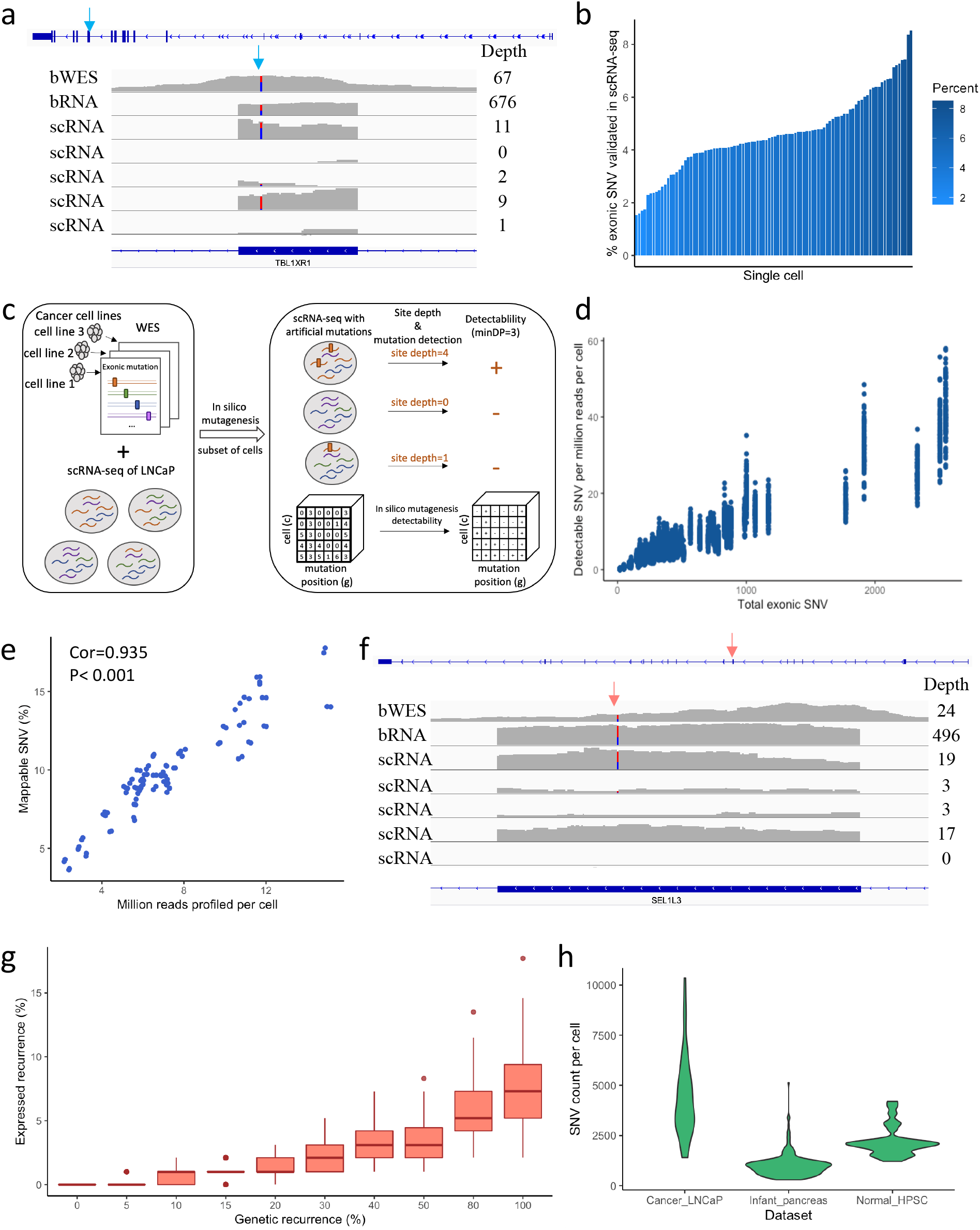
Sparsity and noisiness of expressed somatic mutations in scRNA-seq data. a, Genome browser view of bulk WES (bWES), bulk RNA-seq (bRNA), and scRNA-seq (scRNA) profile for the example of a hotspot mutation in the TBL1XR1 gene exon in the LNCaP cell line. Site-specific depth of the mutation loci is labeled on the right side of the browser view. b, Percent of LNCaP exonic somatic SNVs validated in scRNA-seq data, where single cells are ordered with increasing coverage. c, Workflow of the *in silico* mutagenesis for 100 sets of cancer exonic somatic mutations using the LNCaP single cell transcriptomic background. d, Positive correlation between the number of simulated exonic SNV and detectable SNV per million reads per cell. e, Positive correlation between sequencing coverage per cell and percentage of detectable SNV in scRNA-seq. f, Genome browser view of bulk WES, bulk RNA-seq, and scRNA-seq profile for the example of a somatic SNV in SEL1L3 gene exon in the LNCaP cell line. g, Expressed recurrence as detectable in the LNCaP single cell transcriptomic background with different simulated underlying genetic recurrence in the population. h. High numbers of expressed somatic SNVs per cell detected in three scRNA-seq datasets.

Next, we reasoned that additional factors like mutation burden of the cell, sequencing coverage, and minimum site-specific depth may all affect detectability of somatic variants in scRNA-seq. To better understand how different factor affect detectability, we simulated the detectability of mutation sets from 100 cancer cell lines using *in silico* mutagenesis (Fig. 1c, Methods). We observed that both mutation burden, reflected by total exonic SNV, and single-cell sequencing coverage correlate positively with detectability, while site-specific depth cutoff negatively correlates with detectability (Fig. 1d,e, Supplementary Fig. 1a,b). In addition, expression variation, or site-specific depth variation within the single cell contributes to the detectability variation. Nonetheless, less than 3% of detection rate, defined as percent exonic SNVs that are expressed at detectable level per million reads per cell, was found for most cells (Supplementary Fig. 1b, c), confirming sparsity of expressed mutations. Thus, identifying expressed somatic mutations from scRNA-seq data can be challenging due to sparsity of the data, especially for cells with low mutation burden and shallow sequencing coverage.

In addition, the simulation above underestimates the sparsity of expressed mutations in scRNA-seq. Because the above simulation imposes mutagenesis in every single cell, whereas cancer somatic mutations are typically present in a subset of the cells in the population (referred to as cross-cell recurrence hereafter). For example, the somatic mutation on SEL1L3 gene exon region in the LNCaP cancer cell line showed a variant allele frequency (VAF) of 0.375 as measured by WES, and higher VAF in bulk RNA-seq and scRNA-seq single cells with detectable mutation. However, 2 out of 5 single cells measured with scRNA-seq do not carry this mutation despite reasonably high depth at the site (Fig. 1f). This example demonstrates the complexity of somatic mutations at single cell level cannot be simply approximated by the VAF parameter. Instead, we propose cross-cell recurrence as an important parameter in studying expressed somatic mutations in scRNA-seq data.

To understand the effect of cross-cell recurrence on detectability, we adjusted the simulation by applying mutagenesis in a clonal manner (Supplementary Fig. 1d, Methods). We observed that when requiring detectable mutations to have the same cross-cell recurrence as introduced in the simulation, detection rate drops dramatically as cross-cell recurrence increases (Supplementary Fig. 1e). This is expected given the known effect of both expression dropout and allelic dropout.

Accordingly, the observed cross-cell recurrence of detectable mutations were lower than the simulated genetic cross-cell recurrence, despite a positive correlation (Fig. 1g). Thus, the clonal effect of somatic mutations as represented by cross-cell recurrence affects mutation detectability in scRNA-seq.

Despite the challenge of sparsity, high levels of noise and artefacts in mutations identified by variant callers from bulk RNA-seq data has been reported (Yizhak *et al*., 2019). Similarly, we identified unreasonably high numbers of candidate somatic mutations in scRNA-seq data of normal human primary hematopoietic stem and progenitor cells (HSPCs) and infant primary pancreas cells (Fig. 1h), for which very few somatic mutations per cell are expected. One possible explanation is that the mutation calling algorithms used were not reliable. However, a recent study evaluated performance of several mutation calling algorithms on scRNA-seq data using simulated scRNA-seq reads. Most of the algorithms tested showed satisfying performance with FDR < 0.05 (Liu *et al*., 2019), suggesting reliability of these algorithms in detecting mutations carried by the sequencing reads. This aligns with our knowledge that the experimental process of scRNA-seq, e.g. cell lysis, cDNA conversion, library preparation, etc (Supplementary Fig. 1f) may create artificial variants vastly exceeding the number of true variants. Subsequently, direct application of such mutation calling algorithms on real scRNA-seq data may mislead biological interpretations of the data. Therefore, computational methods that can effectively filter out such experimental noise and artifacts are essential to somatic mutational analysis of scRNA-seq.

### RESA workflow

We present a computational framework RESA, which detects somatic mutation with high precision directly from scRNA-seq data (Fig. 2a-c). The key assumption of this method is that cancer cells evolve in a clonal manner and thus expressed somatic mutations have cross-cell recurrence, whereas the noise and artefacts are likely distributed randomly with small probability of recurrence across the cell population. RESA is composed of three main steps: initial variant calling, filtering and labeling, and modeling and refinement.

**Figure 2.**
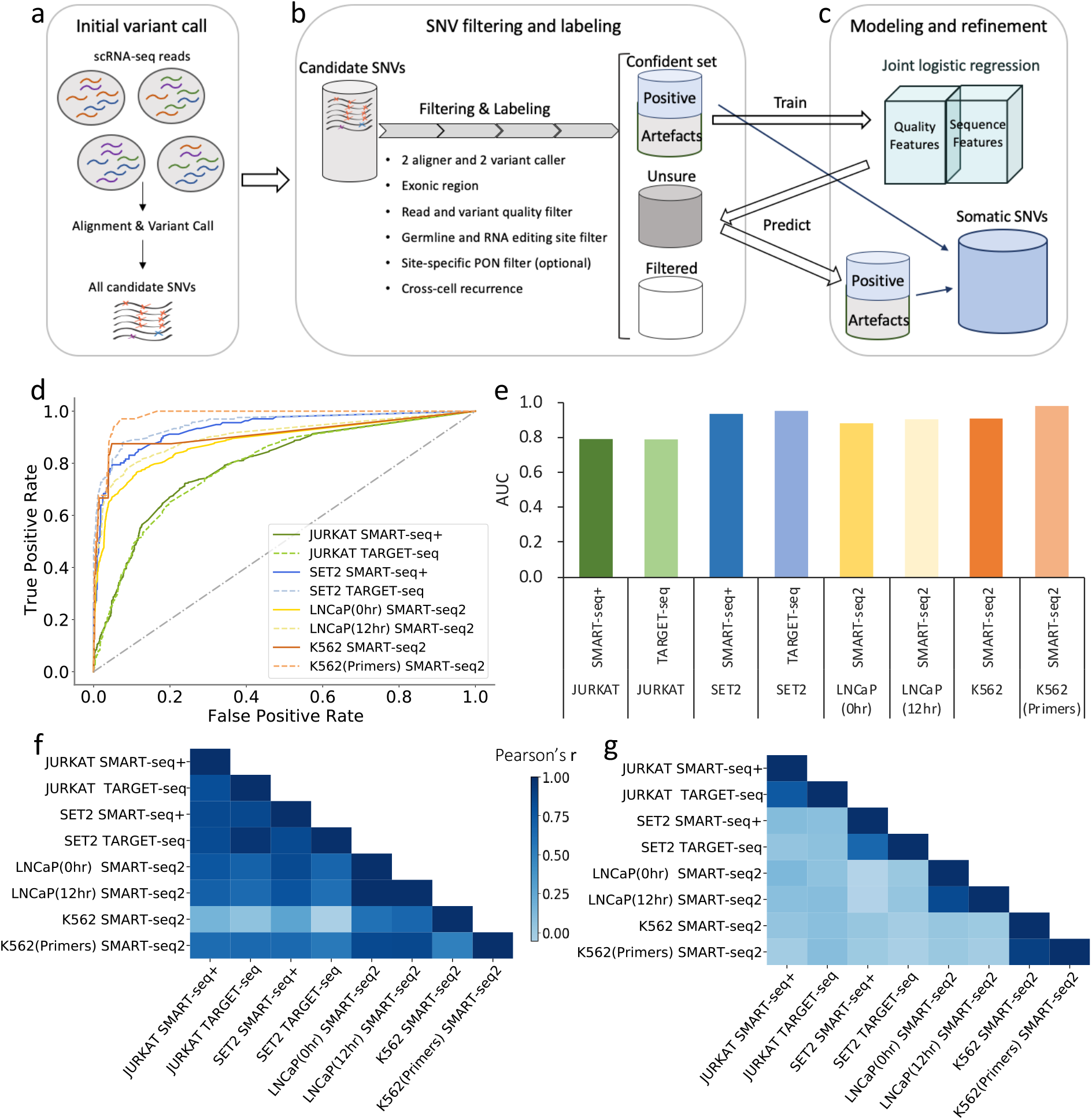
RESA workflow. a, Step1 is initial variant call using two aligner and two mutation calling algorithms. b, Variants called then go through a series of filtering and labeling, categorized into a confident set of positive somatic variants and artifacts, a set of unsure SNVs to refine on, and the rest are filtered out. c, The confident set of variants are used to build a joint logistic regression model, where the model is applied to make predictions in the unsure set of SNVs to refine and expand the final positive set of somatic SNVs. d, the ROC curve, and e, AUC values of the joint logistic regression model prediction on the test set in 8 datasets tested. f, High correlation of feature weights across most datasets observed in quality based logistic regression. g, Cell type specific correlation of feature weights observed in sequence based logistic regression.

The first step of RESA is to align the sequencing reads to the genome and run through mutation calling algorithms with RNA-seq specific parameter settings (Fig. 2a). To minimize aligner and variant caller bias, we apply two independent alignment as well as two variant calling methods. Next, we run through a series of processing, filtering, and most importantly, cross-cell recurrence counting. In this step, the variants are labeled into 3 groups: a high confidence set of SNVs that are putative somatic SNVs or artefact, a set of variants filtered out, and a set of undefined SNVs for further refinement (Fig. 2b, Methods).

Of note, variants that go through the processing and filtering steps without cross-cell recurrence requirement are comparable, if not more strict, to quality based filtering approaches used in other studies (Enge *et al*., 2017; Poirion *et al*., 2018; Liu *et al*., 2019), and are hereafter referred to as hqFilter approach for performance comparison later.

Using the high confidence set of putative positive somatic variants as “positive”, the putative biases and artefacts variants as “negative”, RESA splits the data into training and test sets. We design a joint logistic regression classifier that models both quality and sequence related features independently (Fig. 2c, Methods). The joint logistic regression classifier is then applied to make predictions on the set of undefined SNVs. The final pool of somatic SNVs identified by RESA combines both the positive SNVs from the high confidence set, as well as the predicted positive SNVs from the undefined set using the joint logistic regression model.

We applied RESA to full-length scRNA-seq data of 349 cells from 8 datasets, covering 4 different cancer cell lines generated by different groups, using different experimental protocols, and with different conditions (Methods, Supplementary Table 1). To evaluate the reliability of the joint logistic regression model, we calculated the model performance in the independently held out test set of all scRNA-seq datasets, where the models showed AUC range from 0.79 to 0.98, with average AUC of 0.9 (Fig. 2d, e). In addition, when comparing the feature weights of quality related features in the quality based logistic regression, we observed high agreement across most datasets. This indicates the effect of quality related features on mutation detection could be generalizable (Fig. 2f). In contrast, the feature weights of sequence related features were cell type specific, where only features between different experimental conditions of the same cell line showed high correlation (Fig. 2g). This demonstrates the flexibility of RESA to adapt to specific cell types, and robustness of RESA across experimental conditions.

### RESA achieves high precision in identifying somatic SNVs from cancer cell line data

To overcome noisiness of mutations called from scRNA-seq data, previous studies applied different filtering methods (Darmanis *et al*., 2017; Enge *et al*., 2017; Poirion *et al*., 2018; Liu *et al*., 2019; Maynard *et al*., 2020). However, no specific filtering criteria has been agreed on amongst different studies, and no systematic comparison of these methods have been performed. We summarized the commonly adopted filtering approaches into two main categories: hqFilter and Whitelist-stratify. The hqFilter approach is filtering based on read and mutation quality as mentioned above. This approach assumes that the majority of noise and artefact mutations have low quality metrics as presented in read quality score, variant quality score, strand bias, variant positional bias etc. Another widely adopted filtering approach is to filter variants based on predicted functional impact, and stratify from cancer somatic mutations manually curated in COSMIC database, sometimes with additional requirement of tissue specificity (Methods). This approach is presumably stringent and focuses on identifying a whitelist of driver mutations biased by functional prediction and database curation. While intuitively reasonable and widely adopted, the sensitivity, reliability, and precision in particular, of this approach has not been evaluated. We adopted both methods to compare to the RESA approach in somatic SNV detection.

The availability of WES/WGS data of cancer cell lines from CCLE allowed us to check for detectability of true somatic mutations in the scRNA-seq dataset. Additionally, this enabled us to evaluate the performance of different methods using orthogonal data. The precision of RESA validated by WES across the above-mentioned 8 datasets ranged from 0.63 to 0.9, with average precision at 0.71, whereas the average precision of hqFilter approach is 0.064 and Whitelist-stratify approach is only 0.061 (Fig. 3a). Importantly, the high precision of RESA was not a mere effect of aggressive filtering reflected by the number of mutations remaining after filtering. For the Whitelist-stratify approach, precision remained low regardless of the number of mutations after filtering (SupplementaryFig. 2a). This indicates that quality and functional impact may not be the main characteristics differentiating true mutations in the genome from experimental noise and artefacts. More importantly, noise and artefact SNVs in one dataset might “happen to be” curated cancer somatic mutations that were profiled across thousands of samples and individuals.

**Figure 3.**
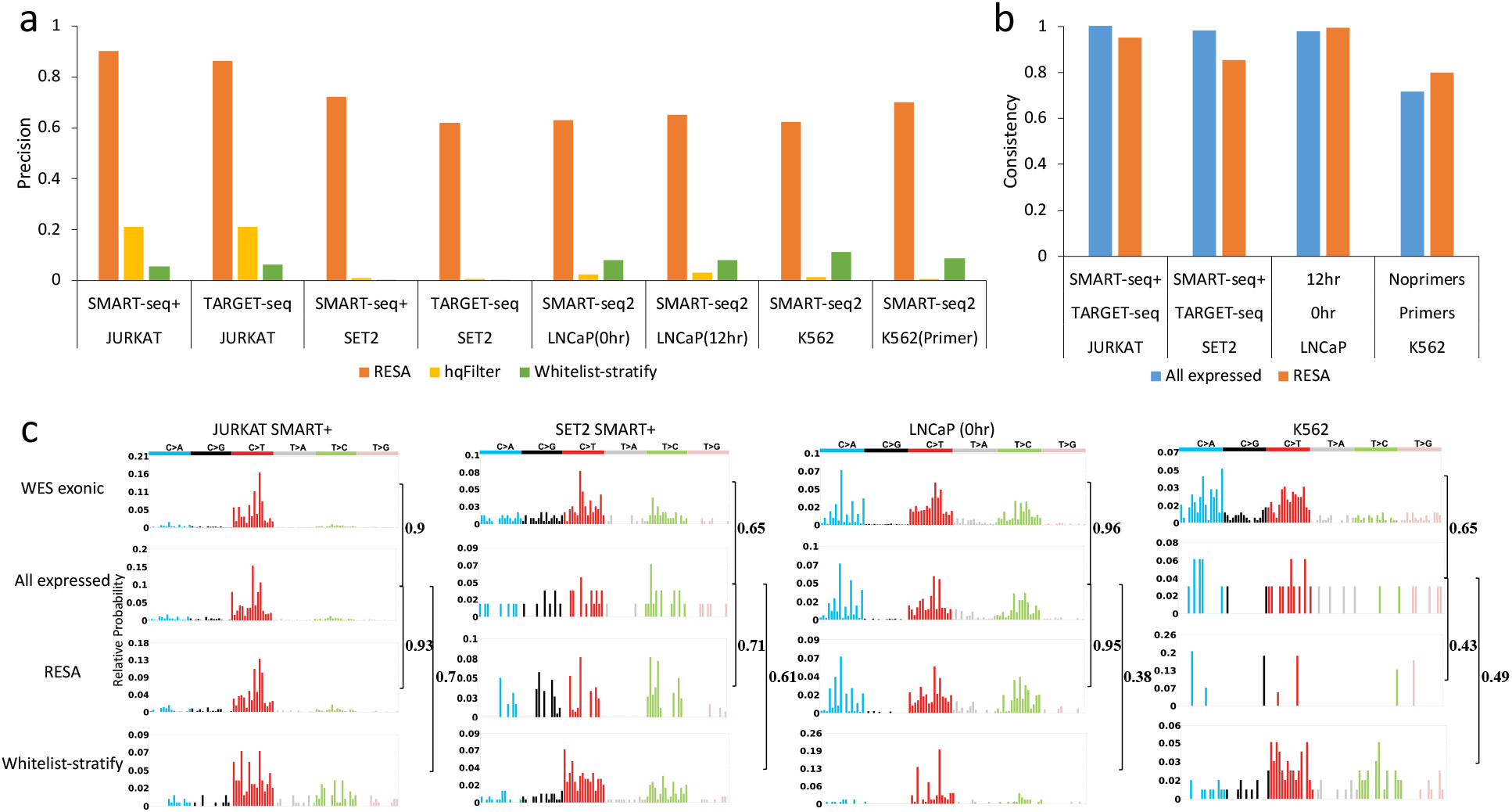
Evaluating precision of RESA with comparison to other methods. a, Barplot showing precision of different methods in identifying positive somatic SNVs using WES data as ground truth across 8 scRNA-seq datasets. b, Barplot showing consistency of different cell line conditions for the same cell line using all the expressed SNVs and RESA. c, Mutation spectra of somatic SNVs identified using all exonic SNVs, all expressed SNVs, RESA, and whitelist-stratify approach across 4 scRNA-seq datasets. Pairwise cosine similarity score was shown next to the brackets.

It is possible that some somatic mutations expressed in a single cell may not be easily detectable through WES/WGS, thus lowering the assessed precision value than the true underlying precision. Clonal divergence between cell cultures that were used to generate WES/WGS vs scRNA-seq data might also lower the precision. However, the fact that RESA reached an average precision of 0.71 across 8 datasets suggested that using expressed exonic mutations detected by WES/WGS is a decent approximation. Of note, comparing expressed somatic mutations of the same cell line under different experimental technologies or conditions, the consistency of all expressed SNVs ranges from 0.71 to 1.02, with an average of 0.93 (Fig. 3b), suggesting that expressed cancer somatic mutations can be replicated reasonably well in independent experiments. Concordantly, the average consistency of RESA is 0.90, again validating the reliability of RESA.

Since SNVs in scRNA-seq data are expected to be sparse, we consider precision as much more important than sensitivity in practice. Regardless, RESA successfully detected somatic SNVs across cancer cells with a wide range of mutation burden, despite variable sensitivity (Supplementary Fig. 2b). In the case of the JURKAT cell line, RESA detected a median of 449 and 387 somatic SNVs per cell, in the SMART-seq+ and TARGET-seq datasets respectively, which was more than 100 times the number of mutations detectable in single cell simultaneous experimental profiling (Supplementary Fig. 2c)(Nam *et al*., 2019; Rodriguez-Meira *et al*., 2019).

In addition to mutation specific evaluation, we also assessed the population-wise mutational spectrum captured by different methods against the WES mutational spectrum, and all expressed SNVs potentially detectable in scRNA-seq data. We observed that expressed SNVs generally captured the exonic mutational spectra well, with lower numbers of SNVs detected and sparse presentation for low mutation burden samples (Fig. 3c). Expressed SNVs detected by RESA faithfully reproduced the mutational spectrum of all expressed SNVs, especially in samples with high mutation burden (Fig. 3c). In contrast, Whitelist-stratify failed to capture the mutational spectra of neither expressed nor all exonic SNVs (Fig. 3c). Thus, expressed SNVs detected by RESA from scRNA-seq data can be used to assess the genomic mutational spectra, especially for samples with high mutation burden.

### RESA reveals complex roles of expressed mutations in melanoma MRD

We applied RESA to a melanoma patient-derived xenograft (PDX) model scRNA-seq dataset, where a time-course study of RAF/MEK inhibition resistance was conducted at four time points in over 600 single cells (Rambow *et al*., 2018). The four time points are T0, phase1, phase2, and phase 3, which corresponds to tumor before treatment, tumor shrinking stage, minimal residual disease (MRD) stage, and relapsed stage after drug treatment, respectively. Reanalysis of the dataset using graph-based clustering of the expression profile identified six clusters of cancer cells (Fig. 4a). Overlaying the four time points to the cluster, we observed clusters unique to, as well as shared across timepoints (Fig. 4b, Supplementary Fig. 3a), confirming the expression heterogeneity in the scRNA-seq of the melanoma samples across different drug treatment stages. For example, cluster 2 and 6 represent groups of cells that are unique in the MRD stage of phase 2, whereas cluster 4 represents cells whose expression patterns are shared by different phases. For the MRD stage of phase 2, cell clusters of 0,1,2,3,4,6 are all present, indicating coexistence of heterogeneous subclones rather than single dominating subclones at MRD despite minimal tumor size.

**Figure 4.**
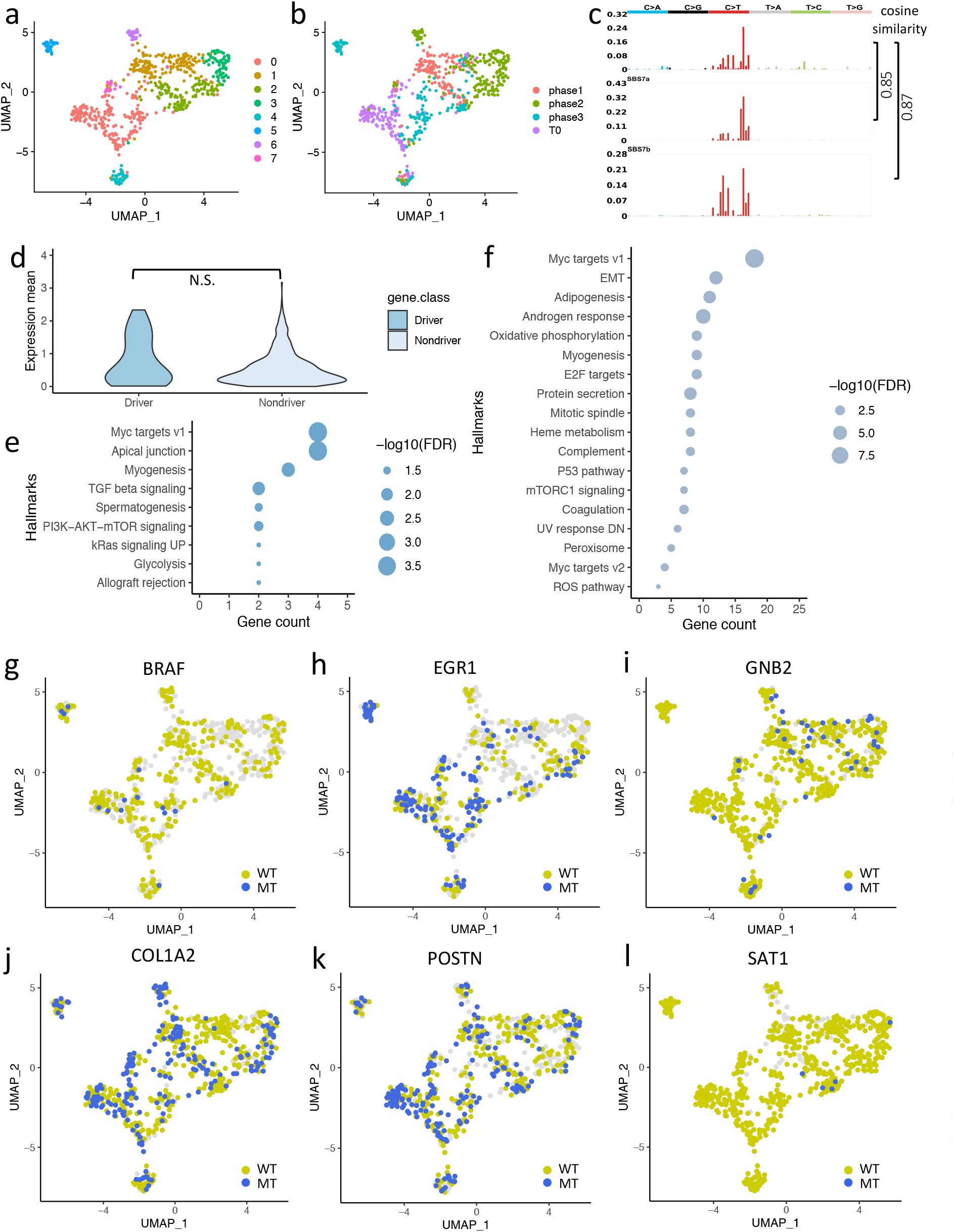
Reanalysis of a melanoma scRNA-seq dataset using RESA. a, Umap plot showing 6 clusters identified by graph-based clustering of the expression profile. b, Umap plot with different color labels for cells from different tumor stages. c, Mutational spectra of somatic variants identified by RESA recapitulates UV dominated signature. Cosine similarity to COSMIC cancer mutation signature 7a and 7b is labeled on the right. d, Violin plot showing no significant difference between the mean expression of driver vs non-driver genes across cell population. e, Cancer hallmarks enriched in driver genes harboring somatic mutations. f, Cancer hallmarks enriched in non-driver genes harboring somatic mutations. g-l, Umap plot comparing distribution of wild-type only (yellow) and mutant state (blue) cells for BRAF (g), EGR1 (h), GNB2 (i), COL1A2 (j), POSTN (k), and SAT1 (l). Cells with undetectable gene of interest is colored in gray.

We successfully identified the BRAF V600E driver mutation of the sample using RESA (Fig. 4g). Though bulk WES/WGS data is not available to quantify precision of RESA, the fact that melanoma tumors are dominated by UV signature in their mutational spectra provides an opportunity to assess the overall reliability of RESA. Applying RESA to this dataset, we successfully recovered the mutational spectrum of typical melanoma samples that are dominated by signature 7, or the UV signature (Fig. 4c). Interestingly, the UV signature remains strong with little variation across different clusters or time points of the cell population, despite expression heterogeneity of somatic SNVs and involved genes (Supplementary Fig. 3b). This is consistent with previous knowledge that treatment with mechanism of action (MOA) not involving DNA replication and repair, e.g. BRAF inhibitor dabrafenib, is not expected to induce large mutational changes (Rambow *et al*., 2018).

The main focus of past somatic SNV cancer studies from scRNA-seq data has been in driver genes, with the assumption that somatic SNVs in driver genes are easily detectable with high expression and biological significance. However, whether expressed somatic SNVs in driver genes are generally expressed at higher levels, or carried in more cells in the population, than those in non-driver genes remain unclear. In this dataset, we identified 575 unique somatic SNVs using RESA, which mapped to 524 genes (Supplementary Table 2). We identified about only 5% or 26 driver genes out of the 524 genes harboring somatic SNVs (Methods, Supplementary Table 3). Comparing driver genes to non-driver genes, no significant difference was found in expression level across cell population (Fig. 4d). This suggests that while driver genes play a key role in cancer initiation and progression, expressed somatic SNVs in non-driver genes can be as detectable as driver genes in scRNA-seq data.

To explore the mutation associated expression signatures, we tested the cancer hallmarks enriched for genes harboring expressed somatic SNVs. In addition to cancer hallmarks shared between driver and non-driver genes, a number of additional cancer hallmarks were found in non-driver genes (Fig4. e,f), suggesting genetic perturbations in many more cancer hallmarks carried by expressed mutations of non-driver genes. Indeed, numerous studies have demonstrated that melanoma development and progression involves p53 pathway, UV response, epithelial to mesenchymal transition(EMT) pathway, androgen response pathway, oxidative phosphorylation pathway, etc (Stretch *et al*., 1991; van Elsas *et al*., 1996; Box, Vukmer and Terzian, 2014; Schmidt *et al*., 2016; Y. Wang *et al*., 2017; Tang *et al*., 2020). Interestingly, when comparing hallmarks of non-driver gene mutations in different stages, DNA damage repair was enriched only in the MRD stage, indicating a potential mechanism of tumor survival in this case (Supplementary Fig. 4). Therefore, hallmarks potentially perturbed by expressed somatic SNVs revealed important orthogonal information in the mechanism of tumor progression on top of expression variation.

To explore the interplay between genomic and transcriptomic heterogeneity, we compared the distribution between mutant containing cells and wild type only cells on the UMAP. We reasoned that mutations that play important roles in specific expression signatures often have elevated expression, thus more likely to be detected by RESA. In addition, such mutations are likely nonuniformly distributed in the cell population compared to its wild type counterparts, and may be enriched in specific tumor stages or expression clusters. Scanning through all the mutations identified by RESA, we found 274 mutations with nonuniform distribution compared to wild type across all the cell population (Adonis test, p<0.05). Furthermore, we identified 7 mutations enriched in the MRD stage of phase 2, and 11 mutations enriched in phase 3 (Fisher’s exact test, Supplementary Table 4).

Next, we took a deeper dive into specific gene examples that harbor expressed somatic SNVs identified by RESA. In concordance with the reported amplification of BRAF in phase 3, the BRAF mutant was significantly enriched in phase 3 compared to BRAF wild type (Fisher’s exact test p=0.02, Methods) (Fig. 4g). Intriguingly, in addition to enrichment in the expression cluster 0 that was shared by T0 and phase 3, BRAF mutant was also enriched in expression cluster 5, a cluster uniquely appearing in phase 3. This indicates that the BRAF mutant not only drives tumor progression in cluster 0, where a relapsed subclone likely evolved from, but also likely drives the unique expression subclone of cluster 5 in phase 3.

Similar to BRAF mutant, ribosomal protein RPL5 and transcription factor EGR1 manifest enriched mutant expression in phase 3 (Fisher’s exact test p=1.6e-5, 3.7e-3, respectively), and the unique subclone of cluster 5 (Fig. 4h, Supplementary Fig. 5a). Both genes are commonly mutated and involved in cancer progression in melanoma (Sells *et al.*, 1995; Schmidt *et al.*, 2016; Fancello *et al.*, 2017; Oršolić *et al.*, 2020). In contrast, mutations of GNB2 and SEMA3C genes were enriched in the MRD stage of phase 2 (Fisher’s exact test p=0.003, 1.9e-4, respectively), but are rather depleted in the relapsed stage of phase 3 (Fig. 4i, Supplementary Fig. 5b). Of note, despite distinct subclonal enrichment, both RPL5 and GNB2 were involved in the androgen response pathway, which is related to melanoma tumor growth and invasion. Figure 4jl again showed examples of such contrasting mutant distribution of 3 genes, COL1A2, POSTN, and SAT1, involved in the EMT pathway (Fig. 4j-l). The EMT pathway was considered the most lethal feature of cancer, yet mutations of different genes in the pathway might be involved in different stages of the melanoma. Taken together, expressed somatic SNVs may perturb cancer pathways and hallmarks in a stage-specific and mutation-specific manner, revealing a complex relationship between expression and mutational intratumor heterogeneity.

## Discussion

Identifying somatic mutations directly from scRNA-seq data has been a known challenge. While many attempts have been made, no computational method has been proven widely applicable. We report a computational framework named RESA, which identifies expressed somatic SNVs with high precision directly from scRNA-seq data. RESA can effectively filter out noise and artefacts generated during the experimental procedure, thus achieving high precision. In addition, RESA applies a joint logistic regression to expand the putative somatic mutations, which helps increase detection sensitivity while maintaining high precision. We tested RESA across datasets both *in vitro* and *in vivo*, and demonstrated the reliability of RESA. Importantly, we showed that RESA exhibits substantially higher precision than the commonly used quality-based and whitelist-based filtering approaches.

Ideally the detected expressed SNVs in scRNA-seq should be assessed by high quality genotype profiling in the same cell. However, single cell whole genome sequencing (scWGS) is known to contain high levels of noise and artefacts (Huang *et al*., 2015), and technologies that simultaneously profile genomic DNA and mRNA of the same cell are also prone to low sensitivity and high noise level (Macaulay *et al*., 2015; Hou *et al*., 2016; Han *et al*., 2018). Due to the above limitations, the commonly used approximation is to overlap expressed SNVs in scRNA-seq with SNVs detected in WES data. The WES data allows us to assess the precision of the RESA approach at population level. In addition to evaluating individual SNVs, RESA also attempts to assess the mutational spectrum of expressed SNVs. High precision detection of somatic SNVs enables reliable mutational signature analysis of the expressed SNVs in scRNA-seq data, thus providing insights into the mechanistic biological processes involved in cancer progression, or revealing potential therapeutic opportunities in the sample of interest.

Detection of expressed SNVs using RESA may suffer from several limitations. RESA mainly works in scRNA-seq data that are sequenced relatively deep (>3-5 million reads per cell in general), or in samples with relatively high mutation burden. Application to other widely used scRNA-seq technologies like 10X genomics and drop-seq will thus be limited by the nature of shallow sequencing and biased gene region coverage. Another potential limitation is that this approach is designed to detect clonal somatic mutations instead of rare mutations in the population. Thus, the limited sensitivity of RESA might potentially limit its application.

The success of RESA in high precision somatic mutation detection from scRNA-seq data highlights the critical importance of cellular recurrence, whereas prioritizing based on functional prediction or stratification from curated whitelist mutations barely helps. This is the first study, to the best of our knowledge, that made direct comparison among the methods emphasizing the above aspects and delivered distinctive results. As more datasets, especially datasets with both scRNA-seq and WES/WEG profiling on the same sample become available, new methods may be developed with better performance. We believe RESA will provide valuable information to facilitate single cell level genotype to phenotype study in the future.

## Methods

### WES, bulk RNA-seq, and scRNA-seq profile of cancer cell lines

The bam files for WES data and bulk RNA-seq data for cancer cell line LNCaP were downloaded through Genomic Data Commons (https://gdc.cancer.gov/) mapped to GRCh38. Genome browser view of selected genomic regions were plotted using IGV version 2.6.3 (Robinson *et al*., 2011).

The exonic somatic mutations for all cancer cell lines tested in this study were processed by the CCLE project and downloaded from CCLE_20Q1_mutations (Ghandi *et al*., 2019). The scRNA-seq for cancer cell lines were downloaded from GSE105451, GSE99795, and GSE76312, respectively.

Specifically, Rodriguez-Meira et al generated scRNA-seq data for leukemia cancer cell line JURKAT and SET2 using SMART-seq+, an optimized experimental procedure based on SMART-seq2, and TARGET-seq, a scRNA-seq technology that detects selected genomic mutations in parallel with unbiased whole-transcriptome analysis (Rodriguez-Meira *et al*., 2019). This dataset not only provided a baseline to test performance of different methods, but also provided an opportunity to set a positive threshold of experimental cross-platform consistency between different scRNA-seq experiments. Similarly, the erythroleukemic cell line K562 dataset was carried out in two experimental condition groups, with one group added DNA primers designed to capture BCR-ABL fusion (Giustacchini *et al*., 2017). The prostate cancer cell line LNCaP were treated with double thymidine for 12 hours to synchronize the population cell cycle, then single cells were collected at 0hr without drug treatment, 12hr with and without drug treatment and scRNA-seq data were generated using SMART-seq2 (Horning *et al*., 2018). We used the 0hr and 12hr untreated scRNA-seq data to evaluate algorithm performance. Because double thymidine treatment induces replication stress and DNA damage response (Matsuno *et al*., 2019), for LNCaP data we focused on high recurrence mutations that are more likely preserving the initial mutation profile of the cell population. Similar to the previous example, comparison of SNVs between scRNA-seq data for LNCaP collected at two time points can be approximated as positive control for experimental variation.

### Mutation detectability simulation

To collect sets of cancer somatic mutations in the exonic region, we randomly selected 100 cancer cell lines from the Cancer Cell Line Encyclopedia (CCLE) project, collected 100 sets of exonic mutations from the selected cell lines, and and compiled the somatic mutations reported by CCLE (Ghandi *et al*., 2019). For scRNA-seq data, we used a prostate cancer cell line LNCaP dataset (0hr time point without drug treatment condition) profiled with SMART-seq2 technology (Horning *et al*., 2018). This scRNA-seq dataset profiled 96 cells with a wide range of sequencing coverage across cells, which allows us to demonstrate the effect of different factors on the detectability of somatic mutations in scRNA-seq. For the first version of the simulation as shown in Fig 1c, for each set of exonic SNVs, we tested the feasibility of *in silico* mutagenesis of each somatic mutation on the RNA-seq reads by counting the site-specific sequencing depth. The underlying assumption is that only sites in the cell tested with sufficient read depth (as a combined result of gene expression level and overall sequencing coverage) are qualified for *in silico* mutagenesis. In the second version of the simulation as shown in Supplementary Figure 1d, by introducing an additional parameter of recurrence (r), we profiled the site-specific depth of randomly selected r cells and summarized for further analysis.

### Mutation detection using RESA and hqFilter

The details of RESA include three main steps:

### Initial read mapping and variant calling

To minimize aligner and variant caller bias, we run through 2 independent alignment and variant calling methods. The first is a widely adopted procedure, where we align the reads using STAR aligner (Dobin *et al*., 2013), process the aligned reads as suggested by GATK (gatk/4.0.0.0) (DePristo *et al*., 2011), and call variants using GATK HaplotypeCaller. The other procedure is aligning reads using Minimap2 aligner(Li, 2018), followed by Strelka (Kim *et al*., 2018) RNA variant calling settings. For this procedure, only variants marked as passing default filtering set by the Strelka algorithm were preserved. We used STAR-GATK procedure as primary reference for downstream analysis because we observed the GATK process does help identify and correct noise and artefacts from the data.

### Filtering and labeling

With candidate variants identified from the primary STAR-GATK pipeline, we first annotate the variant location and kept only SNVs located in the exonic region. Next, we filter out potential germline variants that are reported in population SNP databases including gnomad 3.0 and dbSNP. Potential RNA editing events are also removed for SNVs that overlap with RNAediting databases.

Next, RESA checks concordance between the two aligner and variant calling pipelines. We use variants identified by both pipelines as candidate variant set A and those not validated by the 2nd pipeline as candidate set B. For each set of variants, RESA further processes the variant calling outputs by setting additional requirements on read quality, variant quality, strand bias and variant positional bias. Also, RESA takes overlap variants from the above-mentioned procedures, requiring variants to have minimum read depth of 3 or user-specified site-specific depth.

Importantly, the resulting filtered SNVs for variant set A are more strict, or at least comparable, than quality-based filtering approaches used in other studies (Enge *et al*., 2017; Poirion *et al*., 2018; Liu *et al*., 2019), and are referred to as hqFilter approach for performance comparison.

After quality-based filtering, RESA sets a cross-cell recurrence filter by keeping only SNVs that are detected in at least X number of cells, X (X>=3) by default is no less than 5% of the total cell number and can be adjusted by user. Also, variants identified in cells more than 80% of the total cell number are considered as putative germline variants and thus are filtered out. The filtered SNVs from set A are considered putative true somatic variants. Thus RESA builds a putative “true positive” set of somatic variants directly from scRNA-seq data based on cross-cell recurrence and quality-based filtering.

To build a high confidence set of noise and artifact SNVs, RESA selects SNVs from set B that failed the above-mentioned criteria, such that putative noise and artefacts have the following properties: 1) SNVs are detected through STAR-GATK procedure but not Minimap2-Strelka procedure; 2) SNVs do not pass the quality-based filtering as described above; 3) SNVs do not show any cross-cell recurrence. We ran through this selection criteria for each single cell, and found very few SNV overlap between the positive and negative set after merging SNVs across the cell population, suggesting decent separation between our putative positive and negative set.

### Modeling and refinement

RESA combines all the putative positive and negative sets of somatic SNVs. The input data are randomly split into training and test sets, with 2/3 of the data used as training. We have designed a classifier joint by two logistic regression models on the training set, and assessed the model performance on the test set. On the training set, the number of data per class is imbalanced and we use the random oversampling replicating observations in the minority class to rebalance the dataset. Next, RESA generates two independent logistic regression models, one based on quality-based features including variant quality, read depth, variant allele fraction, normalized probabilities of genotype and allele depth, and the other using sequence-based features including mutation types, sequence contexts, and mutation signature components. We train the joint logistic regression model using the liblinear library where we applied L1 regularization to the quality-based model, and L2 regularization to sequence-based model. Each logistic regression model returns a probability for positive and negative classes. Then we combine these two models into an integrated classifier with the following equation:

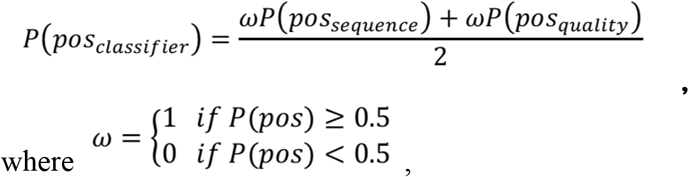

*P*(*pos_sequnce_*) and *P*(*pos_quality_*) are probability of the positive class of the two regression models.

Finally, RESA defines the SNVs positive if *P*(*pos_classifier_*) ≥ 0.5.

After training, we compute the precision, AUC, and accuracy to assess the model performance on the test set. In addition, the model is applied to the candidate SNVs in set B, to extrapolate and refine the putative somatic variant set. This step helped recover some of the true somatic variants that are filtered out in the stringent criteria as described above and thus increases sensitivity while maintaining high precision.

### Mutation detection by Whitelist-stratify

We created a concord whitelist-based mutation selection which has been adopted in several recent studies with similar filtering criteria. Reads were aligned using STAR aligner, processed as suggested by GATK pipeline (gatk 4.0.0.0). Variants were called by GATK HaplotypeCaller (this process is adopted as part of the CTAT mutation caller pipeline) (Fangal, 2020). Next, we filtered out variants using the following criteria: 1) variants are located in exonic region; 2) variant have pathogenic effect as predicted by FATHMM; 3) variant have minimum total read depth of 3; 4) variants have been curated in COSMIC database (Tate *et al*., 2019) in the same tissue type as sample tested.

### Re-analyzing melanoma PDX scRNA-seq expression matrix

The raw reads used for mutation identification and processed gene expression read count table for all cells are downloaded from GSE116237. Expression analysis was done following standard procedure using Seurat package (Satija *et al*., 2015), where read counts are log normalized, with top 2000 variably expressed genes selected for dimension reduction with PCA. Graph based clustering of K-nearest neighbor (KNN) graft following refinement using Louvain algorithm on the top 30 transformed PCs from PCA are applied. UMAP on the top30 PCs with dimension of 20 was used for visualization and visual exploration of the data.

To compare the distribution of mutant containing cells to wild type only cells on the UMAP, for each mutant containing gene, we calculated the Bray-Curtis Dissimilarity between wild type and mutant, and applied Adonis test to calculate p-value. To check for enrichment of mutant containing cells in a specific tumor stage, we compared the cells containing mutant vs wild type only in the stage of interest and the rest of the cell population, and applied one-sided Fisher’s exact test to calculate p-value.

### Driver gene and non-driver gene carrying expressed somatic mutations

We identified mutated driver genes in this dataset by overlapping genes containing RESA identified somatic mutations with COSMIC Cancer Gene Census (CGC) Tier1 gene set, a curated list of genes with validated oncogenic function and was used as reference driver gene set (https://cancer.sanger.ac.uk/census). The rest of the RESA identified mutations containing genes are automatically labeled as non-driver genes.

### Gene-set enrichment analysis

MSigDB hallmarks enriched for driver vs non-driver mutation containing genes were identified from GSEA software (Subramanian *et al*., 2005) with default parameters (https://www.gsea-msigdb.org/gsea/index.jsp). Gene sets corresponding to MSigDB hallmarks were downloaded from https://www.gsea-msigdb.org/gsea/msigdb/index.jsp.

### Custom analysis and plots

Mutation Signature plots were done using SigProfilerPloting. UMAP plots were done using the Seurat package in R. Other custom plots were done using the Seaborn package in python and the ggplot2 package in R.

## Code availability

Source code for RESA pipeline is available on github.

## Acknowledgement

The authors thank Prof. Peter Park, Dr. Doga C. Gulhan, Dr. Simon Chong Chu, and Vinay V. Viswanadham for helpful discussion and suggestions. We thank Dr. Jiaming Li, Dr. Xudong Fu, and Prof. Ruibin Xi for critical reading and suggestions on the manuscript. This work was funded by the Starting Fund from Zhejiang University.

## Author contributions

N.S. designed the project and overall method. N.S. and T.Z. developed, implemented the method and analyzed the data. N.S. drafted the manuscript.

## Reference

Box, N. F., Vukmer, T. O. and Terzian, T. (2014) ‘Targeting p53 in melanoma’, Pigment cell & melanoma research, pp. 8–10.

Campbell, K. R. et al. (2019) ‘clonealign: statistical integration of independent single-cell RNA and DNA sequencing data from human cancers’, Genome biology, 20(1), p. 54.

Darmanis, S. et al. (2017) ‘Single-Cell RNA-Seq Analysis of Infiltrating Neoplastic Cells at the Migrating Front of Human Glioblastoma’, Cell reports, 21(5), pp. 1399–1410.

DePristo, M. A. et al. (2011) ‘A framework for variation discovery and genotyping using next-generation DNA sequencing data’, Nature genetics, 43(5), pp. 491–498.

Dobin, A. et al. (2013) ‘STAR: ultrafast universal RNA-seq aligner’, Bioinformatics, 29(1), pp. 15–21.

van Elsas, A. et al. (1996) ‘Relevance of ultraviolet-induced N-ras oncogene point mutations in development of primary human cutaneous melanoma’, The American journal of pathology, 149(3), pp. 883–893.

Enge, M. et al. (2017) ‘Single-Cell Analysis of Human Pancreas Reveals Transcriptional Signatures of Aging and Somatic Mutation Patterns’, Cell, 171(2), pp. 321–330.e14.

Fancello, L. et al. (2017) ‘The ribosomal protein gene RPL5 is a haploinsufficient tumor suppressor in multiple cancer types’, Oncotarget, 8(9), pp. 14462–14478.

Fangal, V. D. (2020) ‘CTAT Mutations: A Machine Learning Based RNA-Seq Variant Calling Pipeline Incorporating Variant Annotation, Prioritization, and Visualization’. Available at: https://dash.harvard.edu/handle/1/37365605.

van Galen, P. et al. (2019) ‘Single-Cell RNA-Seq Reveals AML Hierarchies Relevant to Disease Progression and Immunity’, Cell, 176(6), pp. 1265–1281.e24.

Ghandi, M. et al. (2019) ‘Next-generation characterization of the Cancer Cell Line Encyclopedia’, Nature, 569(7757), pp. 503–508.

Giustacchini, A. et al. (2017) ‘Single-cell transcriptomics uncovers distinct molecular signatures of stem cells in chronic myeloid leukemia’, Nature medicine, 23(6), pp. 692–702.

Han, K. Y. et al. (2018) ‘SIDR: simultaneous isolation and parallel sequencing of genomic DNA and total RNA from single cells’, Genome research, 28(1), pp. 75–87.

Horning, A. M. et al. (2018) ‘Single-Cell RNA-seq Reveals a Subpopulation of Prostate Cancer Cells with Enhanced Cell-Cycle–Related Transcription and Attenuated Androgen Response’, Cancer research, 78(4), pp. 853–864.

Hou, Y. et al. (2016) ‘Single-cell triple omics sequencing reveals genetic, epigenetic, and transcriptomic heterogeneity in hepatocellular carcinomas’, Cell research, 26(3), pp. 304–319.

Hovestadt, V. et al. (2019) ‘Resolving medulloblastoma cellular architecture by single-cell genomics’, Nature, 572(7767), pp. 74–79.

Huang, L. et al. (2015) ‘Single-Cell Whole-Genome Amplification and Sequencing: Methodology and Applications’, Annual review of genomics and human genetics, 16, pp. 79–102.

Kim, S. et al. (2018) ‘Strelka2: fast and accurate calling of germline and somatic variants’, Nature methods, 15(8), pp. 591–594.

Li, H. (2018) ‘Minimap2: pairwise alignment for nucleotide sequences’, Bioinformatics, 34(18), pp. 3094–3100.

Liu, F. et al. (2019) ‘Systematic comparative analysis of single-nucleotide variant detection methods from singlecell RNA sequencing data’, Genome biology, 20(1), p. 242.

Macaulay, I. C. et al. (2015) ‘G&T-seq: parallel sequencing of single-cell genomes and transcriptomes’, Nature methods, 12(6), pp. 519–522.

Mathys, H. et al. (2019) ‘Single-cell transcriptomic analysis of Alzheimer’s disease’, Nature, 570(7761), pp. 332–337.

Matsuno, Y. et al. (2019) ‘Replication stress triggers microsatellite destabilization and hypermutation leading to clonal expansion in vitro’, Nature communications, 10(1), p. 3925.

Maynard, A. et al. (2020) ‘Therapy-Induced Evolution of Human Lung Cancer Revealed by Single-Cell RNA Sequencing’, Cell, 182(5), pp. 1232–1251.e22.

Nam, A. S. et al. (2019) ‘Somatic mutations and cell identity linked by Genotyping of Transcriptomes’, Nature, 571(7765), pp. 355–360.

Oršolić, I. et al. (2020) ‘Cancer-associated mutations in the ribosomal protein L5 gene dysregulate the HDM2/p53-mediated ribosome biogenesis checkpoint’, Oncogene, 39(17), pp. 3443–3457.

Petti, A. A. et al. (2019) ‘A general approach for detecting expressed mutations in AML cells using single cell RNA-sequencing’, Nature communications, 10(1), p. 3660.

Picelli, S. et al. (2013) ‘Smart-seq2 for sensitive full-length transcriptome profiling in single cells’, Nature methods, 10(11), pp. 1096–1098.

Poirion, O. et al. (2018) ‘Using single nucleotide variations in single-cell RNA-seq to identify subpopulations and genotype-phenotype linkage’, Nature communications, 9(1), p. 4892.

Rambow, F. et al. (2018) ‘Toward Minimal Residual Disease-Directed Therapy in Melanoma’, Cell, 174(4), pp. 843–855.e19.

Robinson, J. T. et al. (2011) ‘Integrative genomics viewer’, Nature biotechnology, 29(1), pp. 24–26.

Rodriguez-Meira, A. et al. (2019) ‘Unravelling Intratumoral Heterogeneity through High-Sensitivity Single-Cell Mutational Analysis and Parallel RNA Sequencing’, Molecular cell, 73(6), pp. 1292–1305.e8.

Satija, R. et al. (2015) ‘Spatial reconstruction of single-cell gene expression data’, Nature biotechnology, 33(5), pp. 495–502.

Schmidt, K. et al. (2016) ‘The lncRNA SLNCR1 Mediates Melanoma Invasion through a Conserved SRA1-like Region’, Cell reports, 15(9), pp. 2025–2037.

Sells, S. F. et al. (1995) ‘The zinc finger transcription factor EGR-1 impedes interleukin-1-inducible tumor growth arrest’, Molecular and cellular biology, 15(2), pp. 682–692.

Stretch, J. R. et al. (1991) ‘Expression of mutant p53 in melanoma’, Cancer research, 51(21), pp. 5976–5979.

Subramanian, A. et al. (2005) ‘Gene set enrichment analysis: a knowledge-based approach for interpreting genomewide expression profiles’, Proceedings of the National Academy of Sciences of the United States of America, 102(43), pp. 15545–15550.

Tang, Y. et al. (2020) ‘EMT-Inducing Transcription Factors, Drivers of Melanoma Phenotype Switching, and Resistance to Treatment’, Cancers, 12(8). doi: 10.3390/cancers12082154.

Tate, J. G. et al. (2019) ‘COSMIC: the Catalogue Of Somatic Mutations In Cancer’, Nucleic acids research, 47(D1), pp. D941–D947.

Wang, L. et al. (2017) ‘Integrated single-cell genetic and transcriptional analysis suggests novel drivers of chronic lymphocytic leukemia’, Genome research, 27(8), pp. 1300–1311.

Wang, Y. et al. (2017) ‘Androgen receptor promotes melanoma metastasis via altering the miRNA-539-3p/USP13/MITF/AXL signals’, Oncogene, 36(12), pp. 1644–1654.

Yizhak, K. et al. (2019) ‘RNA sequence analysis reveals macroscopic somatic clonal expansion across normal tissues’, Science, 364(6444). doi: 10.1126/science.aaw0726.

